# Curvature-driven feedback on aggregation-diffusion of proteins in lipid bilayers

**DOI:** 10.1101/2021.04.02.438263

**Authors:** Arijit Mahapatra, David Saintillan, Padmini Rangamani

## Abstract

Membrane bending is an extensively studied problem from both modeling and experimental perspectives because of the wide implications of curvature generation in cell biology. Many of the curvature generating aspects in membranes can be attributed to interactions between proteins and membranes. These interactions include protein diffusion and formation of aggregates due to protein-protein interactions in the plane of the membrane. Recently, we developed a model that couples the in-plane flow of lipids and diffusion of proteins with the out-of-plane bending of the membrane. Building on this work, here, we focus on the role of explicit aggregation of proteins on the surface of the membrane in the presence of membrane bending and diffusion. We develop a comprehensive framework that includes lipid flow, membrane bending energy, the entropy of protein distribution, and an explicit aggregation potential and derive the governing equations. We compare this framework to the Cahn-Hillard formalism to predict the regimes in which the proteins form patterns on the membrane. We demonstrate the utility of this model using numerical simulations to predict how aggregation and diffusion, coupled with curvature generation, can alter the landscape of membrane-protein interactions.

## 1 Introduction

Cellular membranes contain a variety of integral and peripheral proteins whose spatial organization has biophysical implications for cellular function [1, 2]. In the plane of the membrane, many of these proteins are known to diffuse [3], induce curvature in the bilayer [4], and aggregate either through protein-specific interactions [5] or due to membrane curvature [6]. Interactions between proteins can also lead to the formation of protein microdomains depending on the strength of interaction forces [6, 7]. The ability of these proteins to induce curvature, coupled with the ability of curvature to influence the lateral diffusion-aggregation dynamics, can result in a feedback loop between membrane curvature and protein density on the surface (Figure 1a) [8–10]. In addition to protein aggregation, in-plane viscous flow of the lipid molecules has been found to dominate some of the phase-transition kinetics of vesicle shapes [11]. Recently, we showed that the interaction of membrane bending, protein diffusion, and lipid flow can lead to an aggregation-like configuration on the membrane [12].

**Figure 1:**
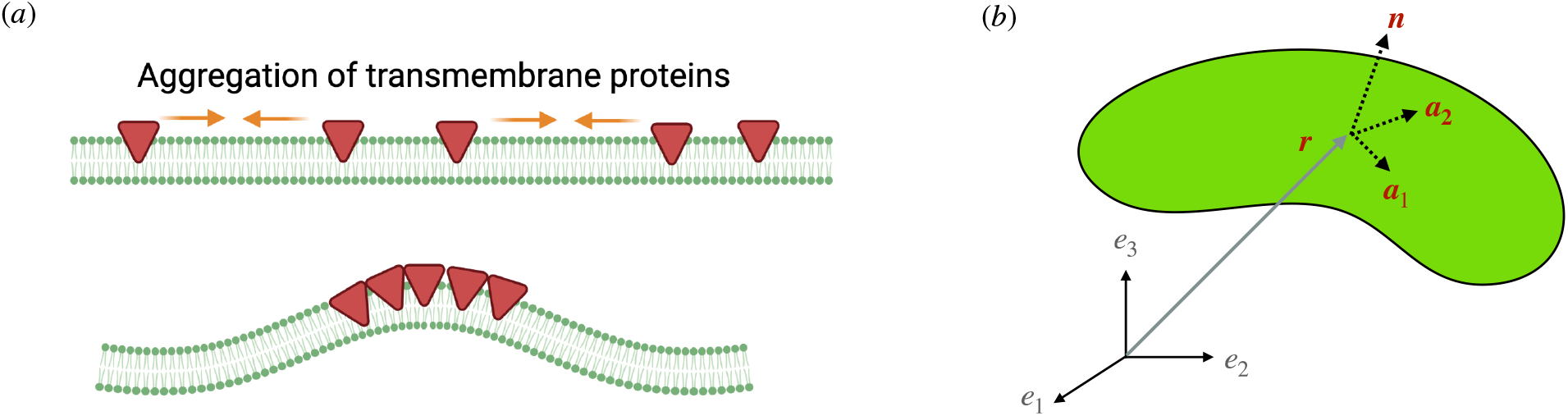
Schematic of protein aggregation and representation of a membrane surface. (*a*) Aggregation of transmembrane proteins on the membrane can lead to domain formation and curvature generation. Here, we develop a continuum model that captures these different interactions. (*b*) Representation of a membrane surface and the surface coordinates. ***a***_1_ and ***a***_2_ are the tangent basis vectors, ***n*** is the unit surface normal.

The aggregation of particles in solvents is a well-studied theoretical problem. Flory [13] and Huggins [14] presented a theoretical formulation for a polymer chain in solution and established the conditions that can lead to its phase separation from the solvent. In binary alloy systems, there has been significant progress on the modeling of the phase transition mechanisms starting from the fundamental Ginzburg-Landau energy [15] that models the interaction energy between the phases as an algebraic expansion in the area fraction of the binary phases around a reference value. Additionally, there are a number of studies that considered the effect of surface tension in the phase separation of solid solutions with an elastic field as a function of concentration field of solute [16–18].

While the classical theories were developed for three-dimensional continua, domain formation and phase separation on two-dimensional surfaces such as lipid bilayers has been of paramount interest recently. The aggregation of proteins on the membrane surface can be viewed as an example of a binary system with lipid and protein as two phases in a two-dimensional curvilinear space. For example, a recent modeling study showed that in a reaction-diffusion system, a pair of activator and inhibitor molecules can lead to an aggregation instability in a specific parameter space, and this instability governs the pattern formation of proteins on membranes [19]. There are many models in the literature that investigate various aspects of phase separation on surfaces. Gera and Salac [10] solved numerically a Cahn-Hilliard system for aggregation-diffusion on a closed torus and observed the temporal evolution of the formation of the aggregation patches. In this case, the surface geometry was fixed. In a subsequent study, they studied the effect of bulk shear flow on the dynamics of the density distribution of species on a deformable vesicle, where the material properties are dependent on the species concentration [20]. Nitschke *et al.* [8] modeled aggregation-diffusion of a two-phase mixture on a spherical surface with in-plane viscous flow, and presented numerical results on pattern formation between the two phases and its strong interplay with the surface flow. The relative interactions between the proteins on the cellular membrane can lead to phase segregation and form protein domains depending on the strength of interaction forces compared to the entropy of mixing [7]. Such aggregation phenomena have been modeled as a polymerization reaction with a very weak free energy of polymerization [7].

Coupling these aggregation phenomena on the membrane surface with membrane deformation is a difficult mathematical and computational problem. Reynwar *et al.* [6] modeled the interaction between proteins with the help of an inter-particle energy and showed that curvature alone can lead to aggregation of these protein particles. A majority of the aggregation studies in the continuum realm consider an aggregation-diffusion chemical potential, which results in the well-known Cahn-Hilliard equation that depicts phase separation dynamics. The energy potential used in studies of protein aggregation on membrane surfaces consists of an inter-molecular aggregation energy and a diffusion potential comprising of the entropy of the protein distribution. Veksler and Gov [21] considered the Ginzburg-Landau energy potential for the aggregation-diffusion energy and modeled the curvature-diffusion instability and identified a parameter space where such instability occurs. Mikucki and Zhou [22] presented a numerical solution for aggregation-diffusion of proteins with bending of the membrane and inviscid flow of lipids. However, their model assumes the local membrane curvature as a function of the density of the proteins as opposed to using a spontaneous curvature, resulting in a weak coupling between bending and diffusion. Givli *et al.* [23] presented a theoretical model of diffusion-aggregation in a multicomponent inviscid stretchable membrane coupled with the bending of the membrane. Additionally, they performed a stability analysis of the system on a sphere, and obtained the most critical modes for the instabilities.

While the models described above capture different aspects of the same problem, here, we sought to develop a comprehensive mathematical model that captures the coupled diffusion and aggregation dynamics, where the proteins induced a curvature resulting in membrane bending and lipids can flow in the plane of the membrane. Such a framework can allow us to explore how the different transport contributions in the plane of the membrane (protein aggregation, protein diffusion, and lipid flow) can contribute both to the formation of protein microdomains and to the curvature generation capability of the membrane. The manuscript is organized as follows. The full system of governing equations is presented in §2. We first analyzed the special case in the absence of bending and reduced the model to a classic Cahn-Hilliard system in §3. We solved the Cahn-Hilliard equation numerically on a square domain and demonstrated the configuration of patch formations in the parameter space. Next, we simulated the fully coupled system in the case of small deformations from a flat plane in §4 and studied the effect of bending energy on the dynamics of aggregation and diffusion of proteins. Our results show that coupling between curvature, protein aggregation, and diffusion can lead to a strong mechanical feedback loop stabilizing the protein microdomains in regions of high curvature.

## 2 Model development

We first formulate the governing equations for coupled diffusion and aggregation of curvature-inducing proteins on a deformable viscous lipid membrane with bending elasticity, building on previous models [12, 24, 25]. We begin by formulating a free energy function for the membrane and apply the principle of energy minimization to derive the governing equations.

### 2.1 Free energy of the membrane

Our system consists of the lipids that comprise the membrane and transmembrane proteins that are embedded in the plane of the membrane and are capable of inducing curvature (Figure 1). Our model does not include the binding or unbinding of proteins from the bulk or the interactions of the bulk fluid with the membrane. The lipid bilayer is modeled as a thin elastic shell, with negligible thickness, that can bend out of the plane and is fluid in-plane. Importantly, we assume that the membrane is areally incompressible and this constraint is imposed on the membrane using a Lagrange multiplier. We describe the different energy contributions to the total energy of the system in detail below.

#### 2.1.1 Protein diffusion

The diffusion of proteins on the membrane surface is modeled using the principle of entropy maximization. The entropy *S* of *q* proteins on *n* binding sites can be found from the multiplicity of the system, Ω = *^n^C_q_*, and is given by

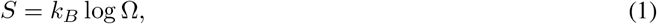

where *k_B_* is the Boltzmann constant. For sufficiently large values of *q* and *n*, the entropic energy per binding site can be represented as a function of area fraction *ϕ* = *q/n* as [26]

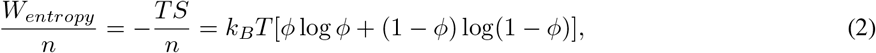

where *T* is the temperature of system. Note that the area fraction *ϕ* can be represented as the ratio of the local protein density *σ* to the saturation density of proteins on the surface *σ_s_*. The energy density per unit area of the membrane is obtained by multiplying the energy density per binding site with the saturation density of the proteins.

#### 2.1.2 Protein aggregation

Aggregation of proteins, on the other hand, can be modeled using the interaction enthalpy of particles in a binary system. With the help of mean-field theory, a continuum representation of the aggregation energy per binding site can be derived as [21, 23, 26]

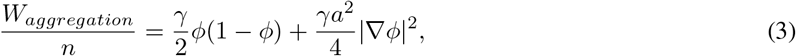

where *γ* is the net effective interaction energy of the proteins of characteristic size *a*.

#### 2.1.3 Bending energy of the membrane

We model the curvature elastic energy density of the membrane using the Helfrich Hamiltonian [27] given by

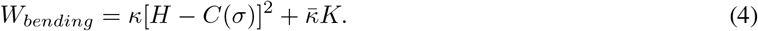

Here, *H* and *K* are mean and Gaussian curvatures of the membrane, *κ* and 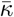 are the bending and Gaussian rigidities, and *C* is the spontaneous curvature induced by the proteins. The spontaneous curvature is assumed to depend linearly on protein density *σ* [12, 25] as

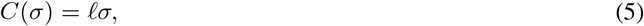

where the proportionality constant ℓ has units of length.

We obtain the total energy density of the membrane, in terms of protein area fraction *ϕ*, by combining Equations (2) to (4) as

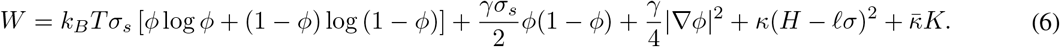

### 2.2 Equations of motion

The lipid bilayer is modeled as a two-dimensional surface, *ω*, in a three-dimensional space (Figure 1b). We refer the reader to [12, 24, 28] for details of the derivation and briefly summarize the key steps here. The surface is parametrized by a position vector ***r***(*θ^α^, t*), where *θ^α^* are the surface coordinates, *α* ∈ {1, 2}. In what follows, (·)_*,α*_ refers to the partial derivative of the quantity in parenthesis with respect to *θ^α^* and (·)_*;α*_ refers to the covariant derivative [29]. The tangent basis vectors are given by ***a**_α_* = ***r***_,α_. The unit surface normal ***n*** is given by ***n*** = (***a***_1_ × ***a***_2_)*/ |**a***_1_ × ***a***_2_|. The metric tensor is given by *a_αβ_* = ***a**_α_ · **a**_β_* and the curvature tensor is defined as *b_αβ_* = ***n · a**_α,β_*. *a^αβ^* is the dual metric and the inverse of *a_αβ_*. *b^αβ^* is defined as *a^αδ^a^μβ^b_δμ_*.

The equations of motions are obtained from a local stress balance on the interface, which can be compactly stated as

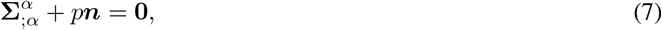

where **Σ** is the stress tensor and 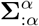 is the divergence of stress. The stress vectors **Σ**^*α*^ can be written in terms of their tangential and normal components as

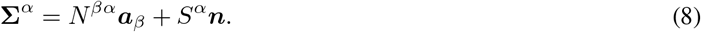

As a result, the local equilibrium of forces, in the tangential and normal directions, is given by [28]

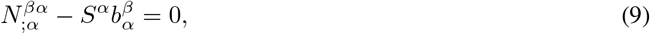

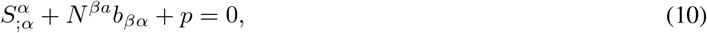

with

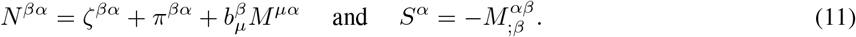

Here, *p* is the normal pressure acting on the surface, *ζ^βα^* and *M^αβ^* are the elastic stress and moment tensors, and *π^βα^* is the viscous stress tensor. The elastic stress and moment tensors can be obtained from the energy density for an incompressible membrane as [12, 28]

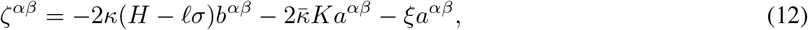

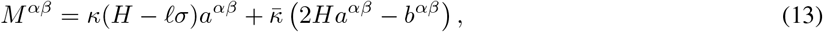

where *ξ* is the Lagrange multiplier that imposes the incompressibility constraint.

The incompressibility constraint on the surface results in the following form of the continuity equation [28]

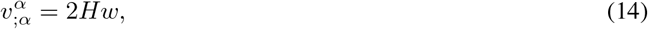

and the viscous stresses obey the constitutive relation [24]

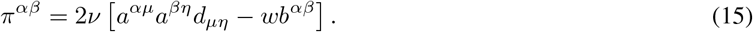

Here, *d_μη_* = (*v_μ;η_* + *v_η;μ_*) /2 is the rate-of-strain tensor expressed in terms of the covariant velocity field *v_μ_* = *a_αμ_v^α^*, and *w* is the normal surface velocity (see [12, 24, 30] for details).

### 2.3 Mass conservation of proteins

Conservation of mass for the protein density *σ* is given by

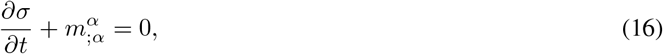

where the flux is

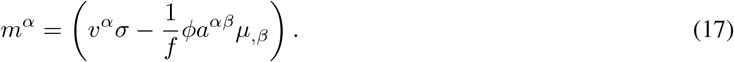

This flux has contributions from advection due to the in-plane velocity field ***v*** and from gradients in the protein chemical potential *μ*. The constant *f* denotes the thermodynamic drag coefficient of a protein and is related to its diffusivity *D* by the Stokes-Einstein relation: *D* = *k_B_T/f*.

The chemical potential, *μ*, is obtained as the variational derivative

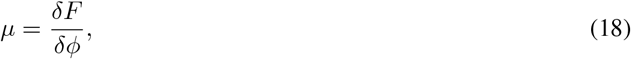

where *F* is the total energy of the system of area *A*:

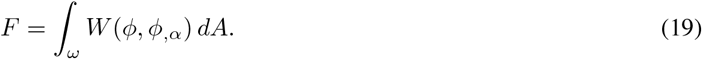

Note that the energy density is a function of both the protein area fraction *ϕ* and its gradient *ϕ_,α_*. Using the definition of the variational derivative, we get the expression of the chemical potential as:

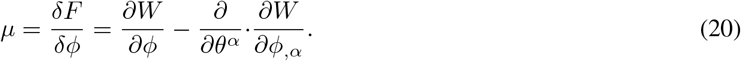

Using Equation (6) for *W*, this yields

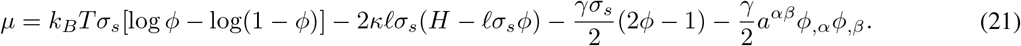

Using Equation (21) in Equation (17) will result in the evolution equation for *σ*.

### 2.4 System of governing equations

Here we summarize the governing equations for the coupled dynamics of the system. Using Equations (11), (12) and (15) for the stresses, the tangential force balance Equation (9) becomes [12, 24, 28]

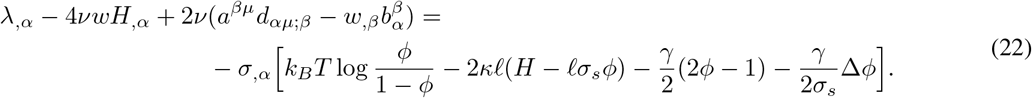

Here we introduce *λ* = −(*W* + *ξ*), which plays the role of surface tension, and Δ(·) = *a^αβ^*(·)_;*αβ*_ is the surface Laplacian. Along with the surface incompressibility condition

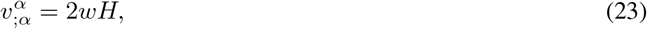

Equation (22) constitutes the governing equation for the velocity field on the evolving surface of the membrane.

The shape of the surface is described by the normal force balance Equation (10), which, after inserting Equation (12), Equations (11) and (15), reads

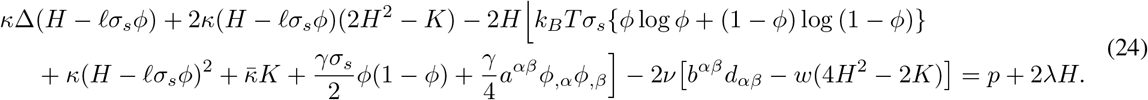

Equation (22) and Equation (24) involve the area fraction of proteins *ϕ* = *σ/σ_s_*, which evolves according to the conservation equation:

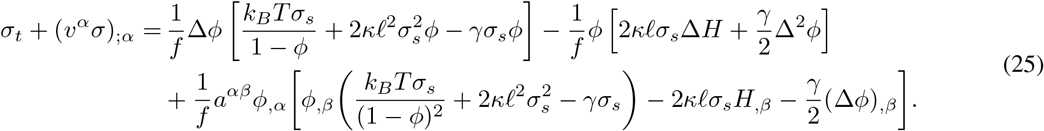

### 2.5 Non-dimensionalization

We non-dimensionalize the system of equations (22)–(25) using the following reference scales. The characteristic length scale is taken to be the size *L* of the domain. The membrane tension *λ* is scaled by its mean value *λ*_0_. Velocities are non-dimensionalized by *v_c_* = *λ*_0_*L/ν*, and we use the diffusive time scale *t_c_* = *L*^2^*/D*. Note that the protein area fraction *ϕ* = *σ/σ_s_* is already dimensionless. Denoting dimensionless quantities with a tilde, the governing equations in dimensionless form are written

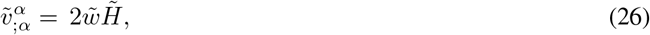

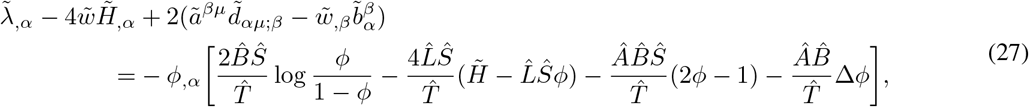

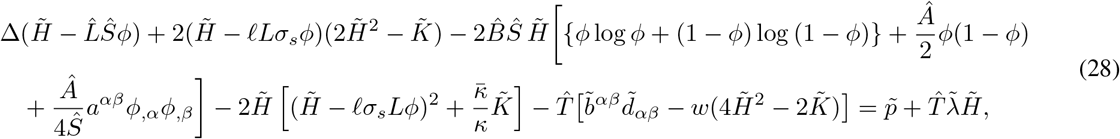

and,

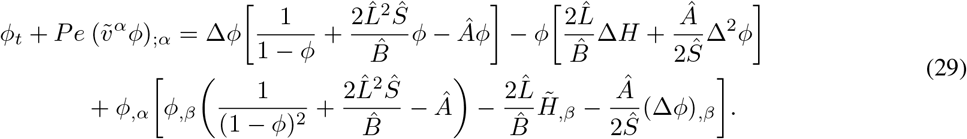

The above system of equations involves seven dimensionless groups that are defined in Table 1 along with their physical interpretation. In all the analyses that follow, we assume that the transmembrane pressure, *p*, is zero. From here on, we also omit tildes on dimensionless variables for brevity.

**Table 1:**
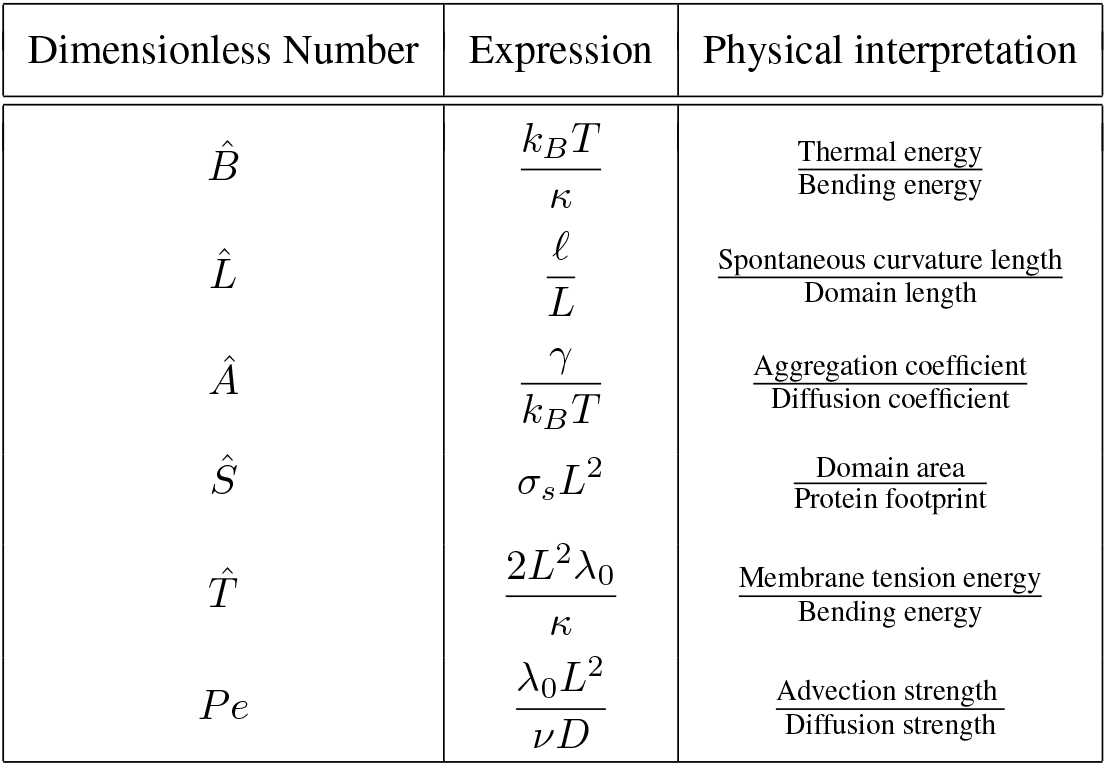
List of dimensionless numbers and their definitions.

| Dimensionless Number | Expression | Physical interpretation |
| --- | --- | --- |
| $\hat{B}$ | $\frac{k_B T}{\kappa}$ | $\frac{\text{Thermal energy}}{\text{Bending energy}}$ |
| $\hat{L}$ | $\frac{\ell}{L}$ | $\frac{\text{Spontaneous curvature length}}{\text{Domain length}}$ |
| $\hat{A}$ | $\frac{\gamma}{k_B T}$ | $\frac{\text{Aggregation coefficient}}{\text{Diffusion coefficient}}$ |
| $\hat{S}$ | $\sigma_s L^2$ | $\frac{\text{Domain area}}{\text{Protein footprint}}$ |
| $\hat{T}$ | $\frac{2L^2 \lambda_0}{\kappa}$ | $\frac{\text{Membrane tension energy}}{\text{Bending energy}}$ |
| $Pe$ | $\frac{\lambda_0 L^2}{\nu D}$ | $\frac{\text{Advection strength}}{\text{Diffusion strength}}$ |

## 3 Cahn-Hilliard system and stability analysis

### 3.1 Reduction to the Cahn-Hilliard system

We first consider the simplified diffusion-aggregation system in the absence of membrane bending and in-plane lipid flow to gain insight into how diffusion and aggregation compete in the plane of the membrane to form protein aggregates (also referred to as patterns or microdomains). We assume that the proteins have zero spontaneous curvature in this case. As a result of these simplifications, the surface gradient (*·*)_*,α*_ is reduced to the planar gradient 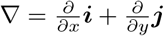 and the surface Laplacian Δ becomes 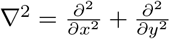. Neglecting the flow and bending terms in Equation (29), we arrive at a transport equation similar to the Cahn-Hilliard equation:

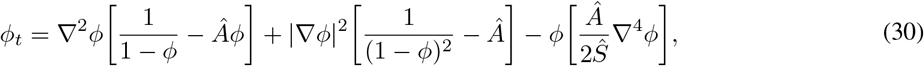

Equation (29) reduces to Fickian diffusion in the dilute limit *ϕ* ≪ 1 in the absence of aggregation (*Â* = 0). Equation (29) is also similar to the system presented by Givli and Bhattyacharya [23], for which they conducted a stability analysis on a closed surface. Here, we present a stability analysis of the equivalent Cahn-Hilliard system on a flat surface, and complement the analysis with numerical simulations of the nonlinear system in a periodic domain.

### 3.2 Linear stability analysis

We perform a linear stability analysis of Equation (30) to identify the parameter regimes that can lead to protein aggregation. The homogeneous state with uniform concentration *ϕ*_0_ is perturbed by a small amount *ϕ*′ such that

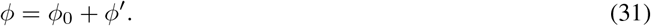

Linearizing Equation (30) provides the equation for density fluctuation *ϕ*′ as

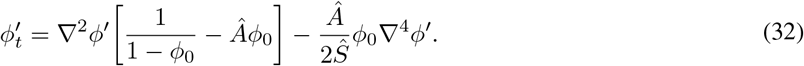

We consider normal modes of the form *ϕ*′ = *e^αt^e^i2π**k**·**x**^*, for which Equation (32) provides the dispersion relation

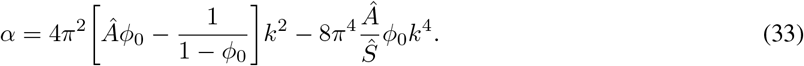

We find that the growth rate *α* is always real. The first term in Equation (33) is positive and therefore destabilizing as long as *Â* ≥ *Â_c_* = [*ϕ*_0_(1 − *ϕ*_0_)]^*−*1^ ≈ 11.1, whereas the second term is always stabilizing. The marginal stability curves *α* = 0 in the (*Â*, *Ŝ*) plane are plotted for various wavenumbers *k* in Figure 2. For a given choice of *Â* and *Ŝ*, this results in a band of unstable wavenumbers 0 *≤ k ≤ k_c_*, where

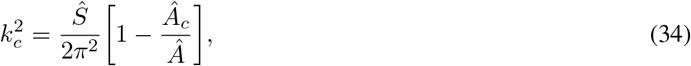

and the maximum growth rate occurs at wavenumber 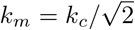. The corresponding wavelength Λ = 2*π/k_m_* provides a prediction for the characteristic lengthscale of aggregation patches, which is expected to decay with increasing *Ŝ* but to increase with increasing *Â*.

**Figure 2:**
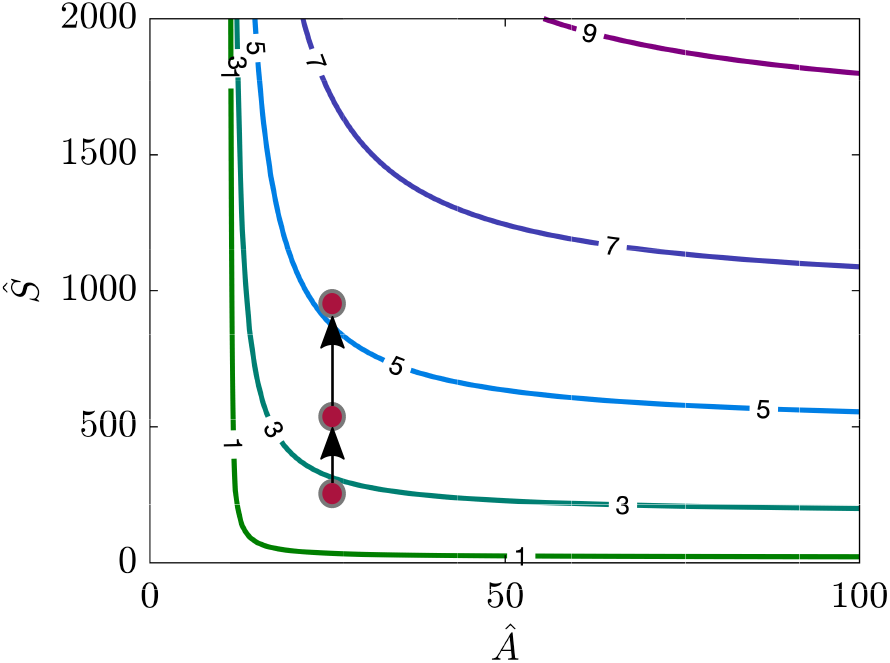
Marginal stability curves for the Cahn-Hilliard system in the (*Â*,*Ŝ*) plane for *ϕ*_0_ = 0.1 and various wavenumbers *k*, as predicted by Equation (30). We mark three points in this figure to identify the parameter values for which we perform nonlinear numerical simulations in Figure 3.

**Figure 3:**
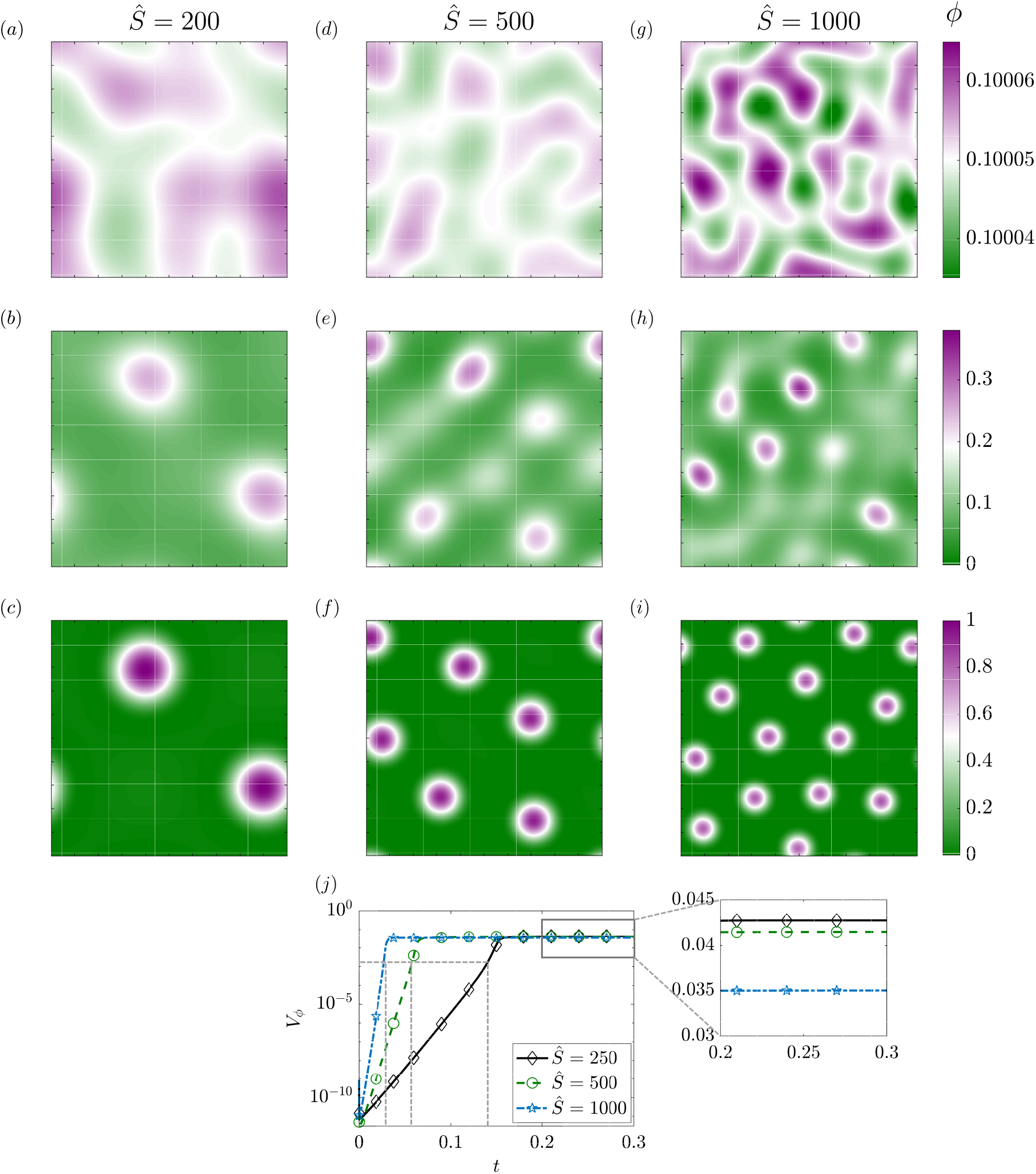
Temporal evolution of the protein distribution in simulations of the Cahn-Hilliard model of Equation (30) on a flat square patch of area 1 *μ*m^2^ for *Â* = 25 and three different values of *Ŝ*. The three rows in panels (*a-i*) correspond to three distinct times: at an early time *t_b_* = 3 *×* 10^*−*3^ shortly after the start of the simulation, at an intermediate time *t_in_* when protein density variance reaches *V_ϕ_* = 2 *×* 10^*−*3^, and at a late time *t_s_* = 0.3 when the system has reached steady state. The three columns correspond to *Ŝ* = 200 (*a-c*), *Ŝ* = 500 (*d-f*), and *Ŝ* = 1000 (*g-i*). Also see Movies M1-M3 in the Supplement for the corresponding dynamics. (*j*) Temporal evolution of the variance *V_ϕ_* of the protein density for the same cases shown in (*a-i*). The dashed lines indicate the intermediate time *t_in_* when the variance reaches *V_ϕ_* = 2 *×* 10^*−*3^.

### 3.3 Numerical simulations

We conducted numerical simulations of Equation (30) inside a square domain for various combinations of *Â* and *Ŝ* that satisfy the necessary condition of aggregation as given in Equation (34) and Figure 2. The initial condition was set as a homogeneous distribution of *ϕ*_0_ = 0.1 with a small random spacial perturbation with magnitude |*ϕ*′| ≤ 1 × 10^*−*4^. We numerically restricted the value of *ϕ* to the interval [*ϵ,* 1 − *ϵ*] with *ϵ* = 1 × 10^*−*3^ to ensure that neither *ϕ* or 1 − *ϕ* becomes zero during the simulations. We used periodic boundary conditions for the protein density and solved the equation numerically using a finite difference technique (the Fortran code is available on Github). In Figure 3, we show the evolution of the protein distribution over time for three different values of the dimensionless number *Ŝ*, while keeping *Â* = 25. In all cases, we find that the initial perturbation in the density field evolves towards the formation of distinct dense circular protein patches that are distributed randomly and nearly uniformly across the domain, in agreement with standard Cahn-Hilliard aggregation dynamics [10]. The main effect of varying *Ŝ*, which is more dramatic than varying *Â* as we further show below, is to control the number of patches as well as their size. Indeed, we recall that *Ŝ*, which is a dimensionless measure of the finite size of the proteins, directly controls the stabilizing term in the dispersion relation Equation (33) and therefore the dominant wavenumber of the instability. Consistent with the stability predictions, we find that larger values of *Ŝ* produce larger numbers of patches with smaller sizes. During the transient evolution, proteins get drawn towards the emerging patches due to aggregation, and at steady state we find that the density inside the patches approached the saturation density (*ϕ* = 1), whereas it approaches zero outside (Also see Movies M1-M3 in the Supplement). We quantify the growth of density fluctuations by plotting in Figure 3*j* the time evolution of the density variance, defined as

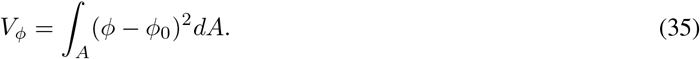

We find that the growth of the variance is exponential at short times, consistent with the expected behavior for a linear instability, before reaching a constant plateau at long times. The growth is observed to increase with *Ŝ* in agreement with the linear prediction of Equation (33). The steady state value, on the other hand, is found to decrease slightly with *Ŝ*, although the differences are small.

A more complete exploration of pattern formation is provided in Figure 4*a*, showing the long-time configurations of aggregated protein patches in the parameter space of *Â* and *Ŝ*. We note that the number of patches, their size, and their homogeneity vary with both parameters. As we already observed in Figure 3, increasing *Ŝ* for a given value of *Â* increases the number of patches and decreases their size. On the other hand, increasing *Â* for a given *Ŝ* tends to increases inhomogeneity among patches, with some visibly denser patches while others tend to be more diffuse. The precise dependence of the number of patches as a function of both *Â* and *Ŝ* is shown in Figure 4*b*, while the steady-state variance is plotted in Figure 4*c*. The variance is found to decrease with *Â*, as the more diffuse patches forming at large *Â* result in weaker spatial fluctuations.

**Figure 4:**
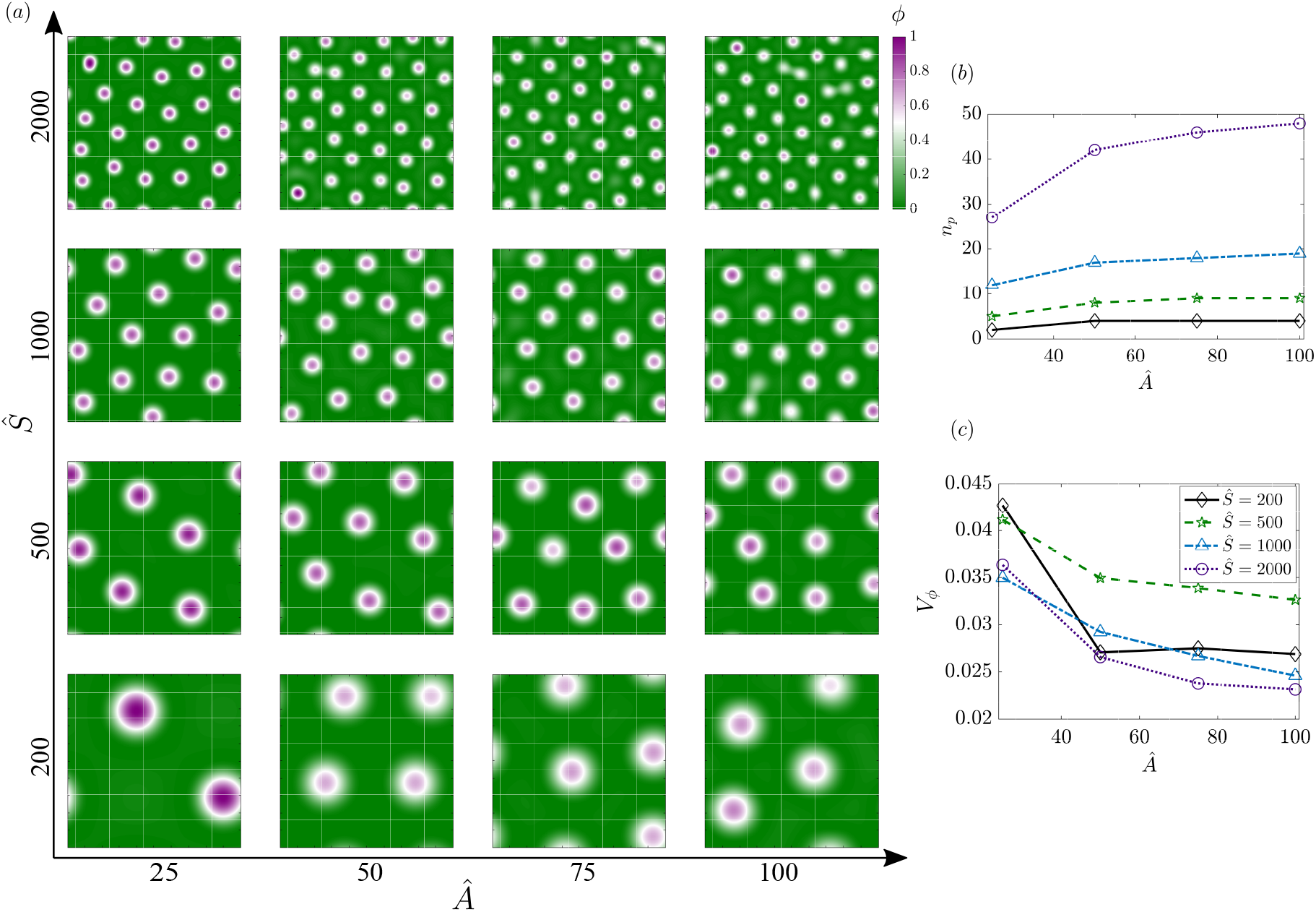
(*a*) Configurations of protein aggregates on a flat square membrane at a late time *t* = 0.3 approaching steady state for various combinations of *Â* and *Ŝ*. (*b*) Variation of the number of protein patches with *Â*, for different values of *Ŝ*. (*c*) Variation of of the protein density variance *V_ϕ_* with *Â* for different values of *Ŝ*.

## 4 Coupling of aggregation with bending: analysis in the small deformation regime

To understand how the inclusion of membrane curvature alters the aggregation-diffusion landscape, we simulated the dynamics of the coupled system Equations (26) to (29) in the regime of small deformations from a plane. The surface is represented using the Monge parametrization, such that the position vector is given by ***r*** = *x_α_**e**_α_* + *z*(*x*_1_*, x*_2_*, t*)***e***_3_. In the regime of small deformations from the plane, we consider gradients of the surface deformation to be small and ignore higher-order terms [12].

### 4.1 Governing equations

The surface gradient and Laplacian in the Monge parameterization are represented as 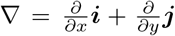 and 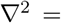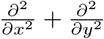. The continuity condition and tangential force balance simplify as

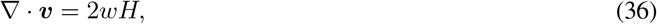

and,

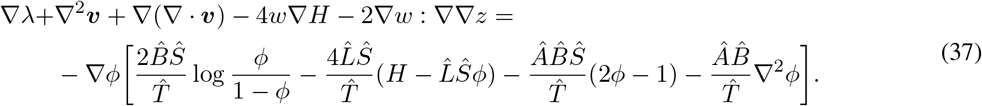

The normal force balance Equation (28) reduces to

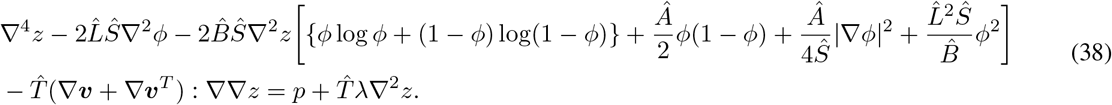

Finally, the transport equation for the protein density field Equation (29) takes on the following form:

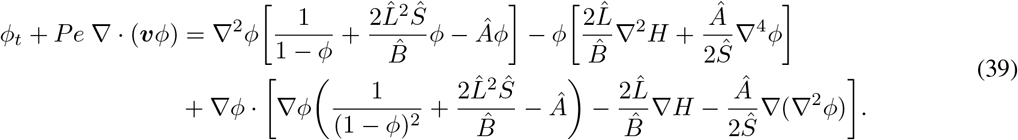

#### 4.1.1 Linear stability analysis

We first perform a stability analysis of the system of equation (Equation (36) to Equation (39)) to identify the parameter regimes similar to Section 3.2, but in the presence of bending due to spontaneous curvature induced by the protein. In the base state, the membrane is flat and at rest with uniform tension (*z*_0_ = 0, ***v***_0_ = **0**, *λ*_0_ = 1), and the protein density is uniform with value *ϕ*_0_. We indeed showed in an earlier study [12] that a uniform protein distribution on a flat membrane is indeed a steady state even when the proteins induce a spontaneous curvature. We then perturbed the variables by small amounts with respect to this base state:

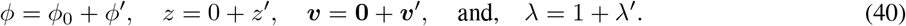

Linearizing Equations (36) and (37) provides the governing equations for velocity and tension fluctuations as

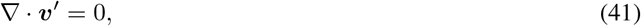

and,

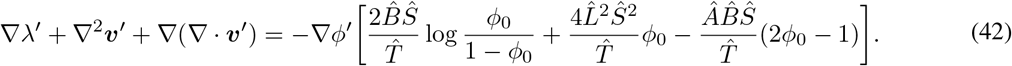

The normal force balance of Equation (38) reduces to

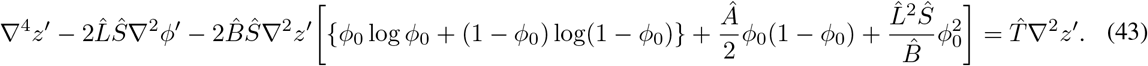

Finally, the transport equation for the protein density given in Equation (39) becomes

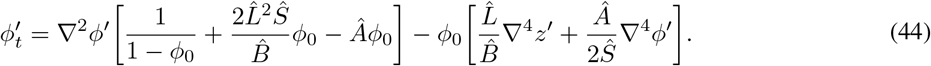

We find that the linearized equations the velocity field and tension completely decouples from the shape equation (43) and protein transport equation (44): in other words, lipid flow and tension fluctuations do not affect the membrane shape and protein transport in the linear regime. To analyze the dynamics of protein aggregation, we therefore need only consider Equations (43) and (44), in which we use the normal modes

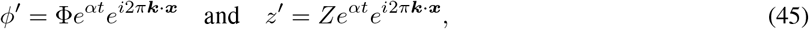

yielding the relations

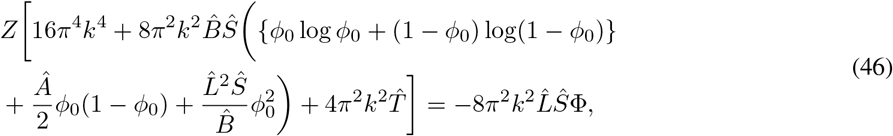

and

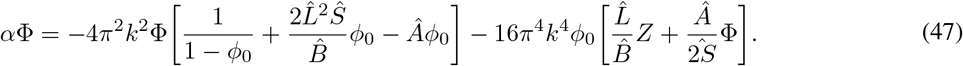

Eliminating variables *Z* and Φ, we obtain the dispersion relation as

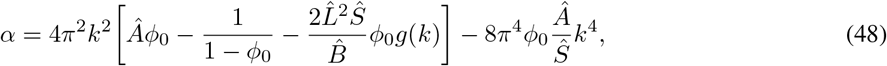

where we have defined

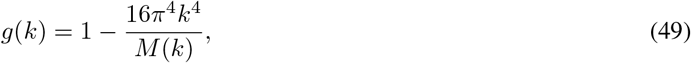

with,

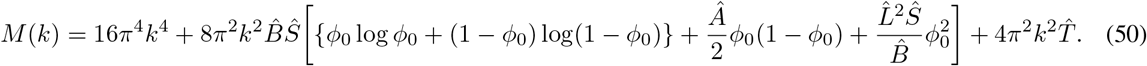

Similar to Equation (33) the second term in Equation (48) is always stabilizing, therefore, protein aggregation happens when first term is positive. The necessary condition for protein aggregates to form becomes

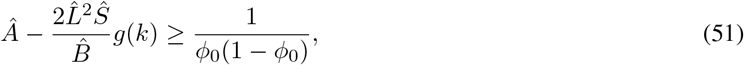

or

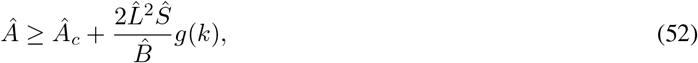

where *Â_c_* is the critical value of *Â* previously introduced in Section 3.2 in the Cahn-Hilliard case. Figure 5 shows the dependence of *g*(*k*) on wave number *k* for *Â* = 25 and various combinations of 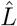 and *Ŝ*. When both 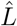 and *Ŝ* increase, *g*(*k*) tends to increases for at small wavenumbers and thus stabilizes the system. This means in particular that proteins with large spontaneous curvature, as captured by 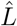, can in fact a stabilizing effect on protein aggregation, and this counterintuitive observation will be confirmed in numerical simulations as we discuss next.

**Figure 5:**
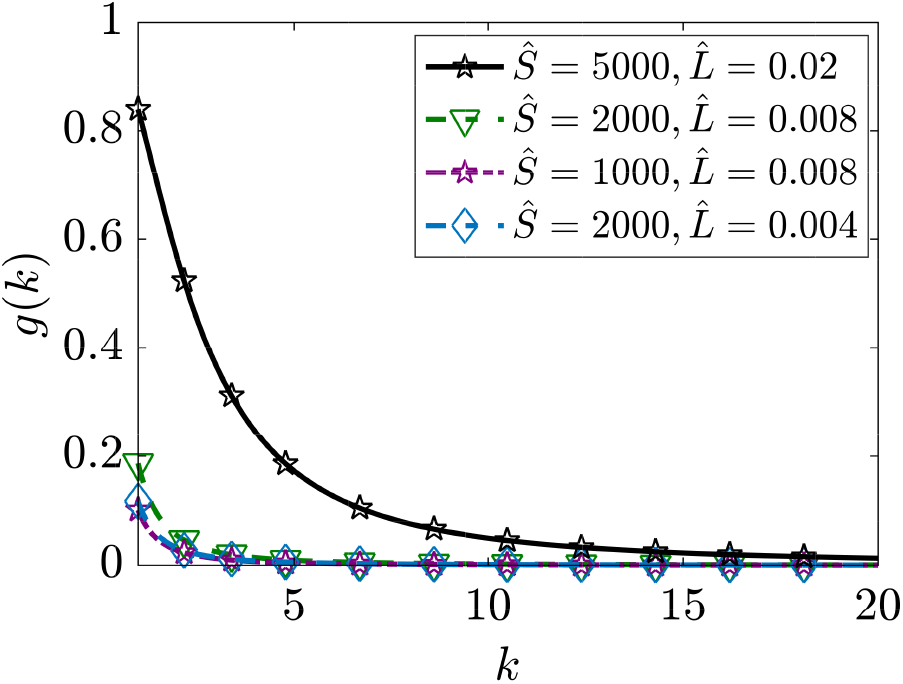
Dependence of *g* defined in Equation (49) on wavenumber *k* for different values of *Ŝ* and 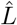, with *Â* = 25.

### 4.2 Numerical Simulations

We solved Equations (36) to (39) numerically on a square domain with periodic boundary conditions for a small random density perturbation over a homogeneous steady state density of *ϕ* = 0.1. The proteins now induce a spontaneous curvature in the membrane, and the model also captures the viscous flow on the membrane manifold. Typical transient dynamics are illustrated in Figure 6 in a simulation with 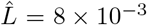, *Â* = 25, and *Ŝ* = 2000. The initial random distribution resolves into strong patches of proteins over time with the same number of patches as we observed in the Cahn-Hilliard system (compare Figure 3*a-c* with Figure 6*a-c*). Because the system of equations now accounts for coupling of curvature with protein dynamics, we observe that the formation of dense protein patches is accompanied by the localized growth of membrane deformations, in the form of nearly circular peaks surrounded by flatter regions of oppositely-signed curvature (Figure 6*a-c*). We also observe that the formation of protein aggregates is coupled with a tangential velocity field in the plane of the membrane, to accommodate the deformation of the membrane (Figure 6*d-f*): as the protein aggregates form and deflect the membrane in the normal direction, a source-like flow is generated locally as dictated by the continuity relation Equation (36). During this process, the magnitude of the velocity increases until the system approaches a steady state where aggregation balances diffusion. As the steady state is approached, the flow in the membrane changes nature as the normal velocity vanishes, with each protein patch driving a weaker flow with quadrupolar symmetry.

**Figure 6:**
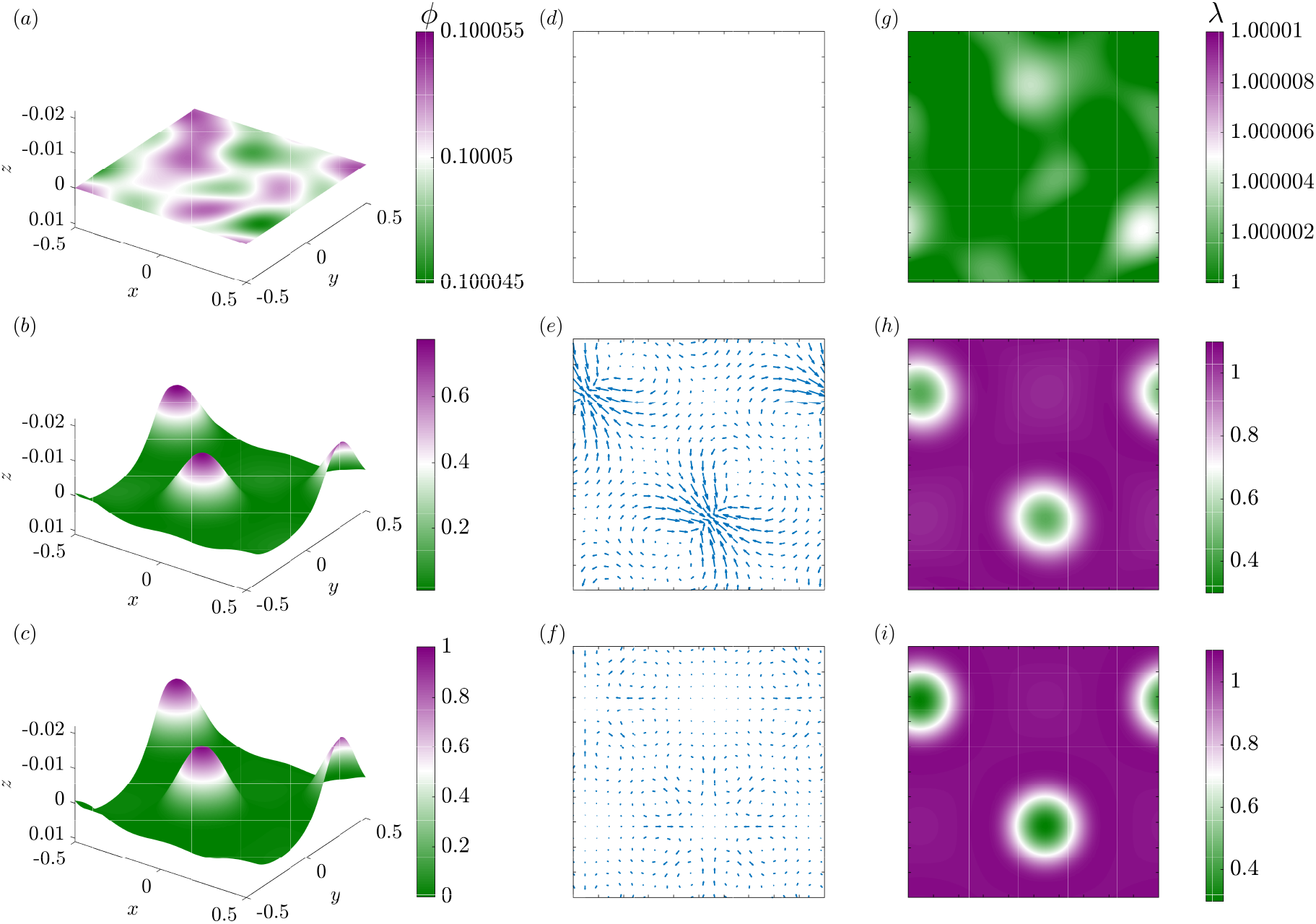
Temporal evolution of protein distribution, membrane shape, in-plane velocity and tension for a square membrane of size 1 *μ*m^2^ with *Â* = 25, *Ŝ* = 200, and 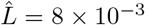. (*a-c*) Height of the membrane colored with the local protein density, (*d-f*) in-plane velocity field, and (*g-i*) membrane tension at dimensionless times 0.003, 0.21, and 0.3.

As we have noted in prior works [12, 24, 30], coupling of lipid flow to membrane deformation not only completes the description of the physics underlying the viscoelastic nature of the membrane but also allows for the accurate calculation of the membrane tension field (the Lagrange multiplier for incompressibility). This is particularly relevant for understanding how microdomains of proteins can alter the tension field in the membrane. The tension field on the membrane tracks with the protein microdomains and the deformation in the coupled system (Figure 6*g-i*). Initially, the membrane has nearly uniform tension, but as regions of high protein aggregation and therefore high membrane curvature form, these locations are found to have lower tension in comparison with the rest of the membrane. Thus, the dynamics of the coupled system is able to capture key experimental observations in the field of membrane-protein interactions: (a) regions of high curvature and aggregation are correlated for curvature-inducing proteins suggesting a positive feedback between these two important factors, (b) lipid flow is important to sustain the deformations, and (c) membrane tension is a heterogeneous field and varies with the local membrane composition.

To further quantify these behaviors, we investigated the parameter space of *Ŝ* and 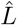 to understand how the spontaneous-curvature induction versus protein footprint compete in a fixed regime of aggregation-to-diffusion (*Â* = 25 fixed) (see Equation (52)). We varied *Ŝ* in the range of 200 to 2000 and 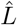 from 1 × 10^*−*3^ to 8 × 10^*−*3^ and summarize these results in Figure 7. We first observed that the growth rate of the variance of *ϕ* shows a strong dependence on 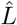 (Figure 7*a*). For *Ŝ* = 200, the growth rate for the two different values of 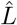 differ slightly with the growth rate being slower for larger 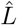. This effect persists and is amplified for larger *Ŝ*: as both *Ŝ* and 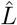 increase, the growth rate decreases, indicating that it takes longer time for patterns to form on the membrane. However, when *Ŝ* = 2000, we see a decay in the variance of protein density *ϕ* as opposed to the exponential growth and eventual plateau for the cases where protein aggregrates form. This result, which is consistent with the stability analysis of Section 4.1.1 suggests that the induction of curvature on the membrane can alter significantly the dynamics of protein aggregation.

**Figure 7:**
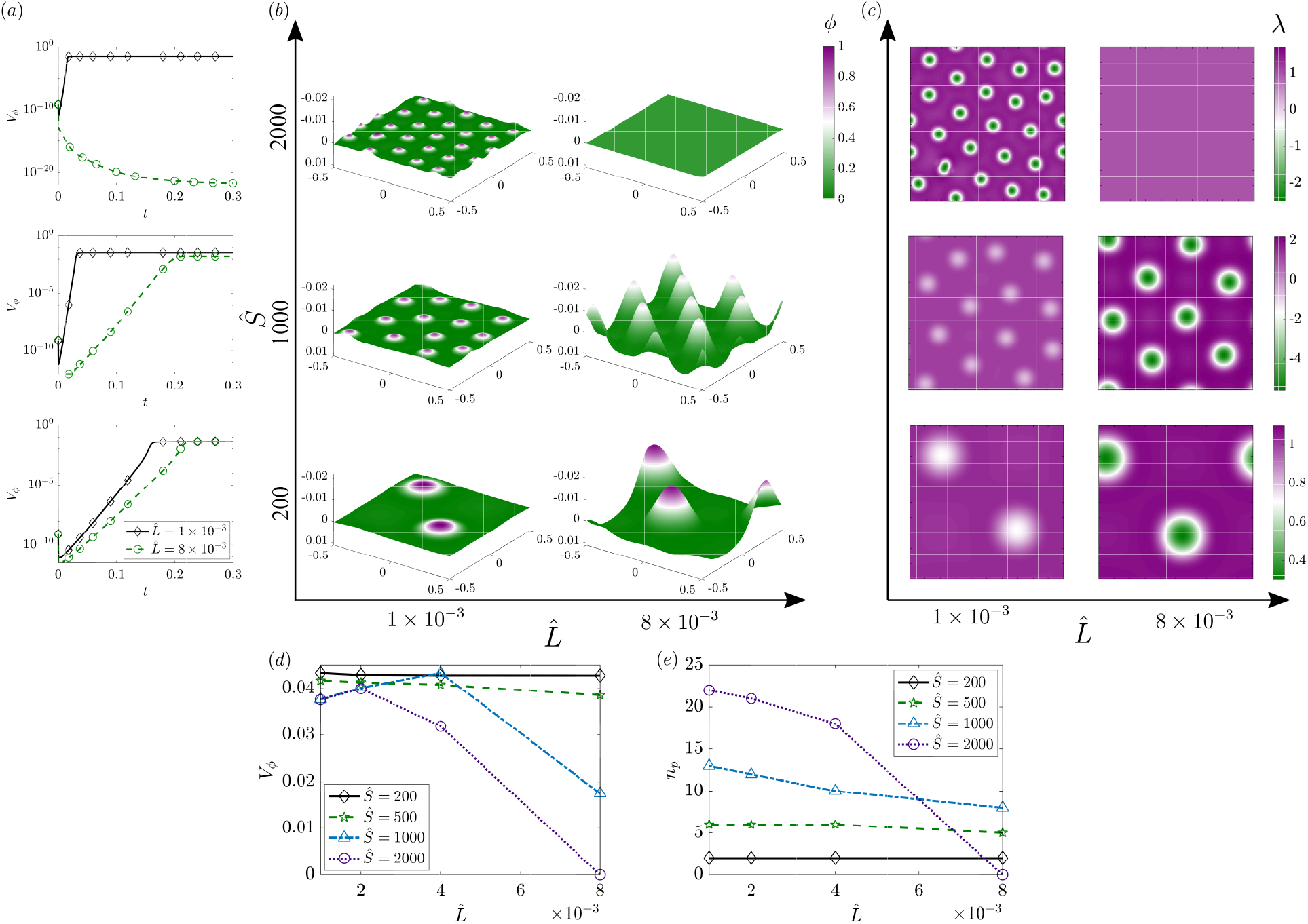
Effect of *Ŝ* and 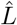 on protein aggregation and membrane dynamics. (*a*) Temporal evolution of the protein density variance *V_ϕ_* for two values 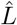 and the same three values of *Ŝ* shown in (*b*). (*b*) Distribution of protein density on the deformed membrane at a long time approaching steady state (*t* = 0.3) for various combinations of 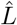 and *Ŝ*, with *Â* = 25. The corresponding dynamics are also shown in movies M4-M6 of the Supplement. (*c*) Distribution of the local membrane tension for the same cases as in (*b*). (*d*) Variance of protein density *V_ϕ_* and (*e*) number of protein patches *n_p_* at *t* = 0.3 as functions of 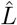, for various values of *Ŝ*.

The steady-state patterns and deformations are illustrated in Figure 7*b* (also see Movies M4-M6, and Figures A.1 and A.2 in the Appendix), where we observe that the number of protein patches is largely unaffected by 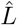 for *Ŝ* = 200. The number of patches increases with *Ŝ* for a given 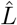 (as already found in Figure 4). However, when *Ŝ* increases to 1000, the number of patches decrease with 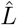. Since the deformation is directly affected by spontaneous curvature, we find however that 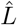 has a strong effect on the magnitude of membrane deflections, with larger protein footprints resulting on stronger deflections. Surprisingly, when *Ŝ* = 2000, we noticed that protein aggregates do not form for the value of 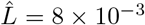 and the membrane remains flat. This phenomenon can be explained from the critical value of *Â* in Equation (52). Since both 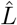 and *Ŝ* have stabilizing effect on density fluctuations *ϕ*′ (Equation (52)), for higher value of *Ŝ* and 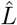, an aggregation coefficient of *Â* = 25 is not sufficient to overcome the stabilizing barrier. However, for lower values of 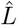 or lower values of *Ŝ*, where the stabilizing effect is relatively weak, we see the formation of protein aggregates.

The tension profile in the membrane follows the inhomogeneity of the protein distribution as expected (Figure 7*c* and Figure A.2). As previously noted in Figure 6, the patches are associated with tension minima. We find, however, that the range of *λ* depends strongly on *Ŝ* and 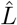, as expected since *λ* is directly proportional to the gradient of the spontaneous curvature, which in turn depends on both ℓ and *σ*. This further highlights the coupling between curvature, flow, and aggregation dynamics. Finally, we look at the variance and the number of patches as a function of both *Ŝ* and 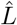 (Figure 7*d,e*). We note that for a given value of *Ŝ*, the variance decreases with increasing 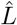 for higher values of *Ŝ* and this decrease is more dramatic when compared to the Cahn-Hilliard model (Figure 4*b*). Even though the number of patches remains more or less unaltered for small values of *Ŝ* as 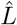 increases, the number decreases with increasing 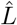 for larger values of *Ŝ* (Figure 7*e*), consistent with the stability behavior noted in Equation (52). These results suggest that the landscape of protein inhomogeneity is not only governed by the *Â*-*Ŝ* space as is the case in the Cahn-Hilliard model; rather the curvature parameters, specifically 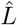 in this case, can have a significant impact on the protein aggregation behavior. Thus, we find that the aggregation-diffusion landscape on the surface of the membrane is altered by the protein-induced spontaneous curvature – tuning these different effects can allow for differential control of curvature-aggregation feedback.

## 5 Discussion

The interaction of peripheral and integral membrane proteins with the lipid bilayer of cellular membranes is fundamental to cellular function [31–33]. In this work, we have developed a comprehensive modeling framework that couples the multiple effects that take place in such membrane-protein interactions: protein diffusion in the plane of the membrane, interaction between the proteins resulting in aggregation, lipid flow in the plane of the membrane, and out-of-plane curvature generation due to protein-induced spontaneous curvature. The resulting system of equations now completely describes the mechanics of a lipid membrane with a second species that can both diffuse and aggregate in the plane of the membrane. We compared this system against a reduced system of Cahn-Hilliard equations to show how the coupling with membrane bending alters the system behavior using both linear stability analysis and numerical simulations. In the absence of curvature coupling (the Cahn-Hilliard system), the dynamics of protein aggregation is driven by the competition between two key parameters, *Ŝ*, representing the relative size of the protein footprint and *Â*, representing the relative strength of protein aggregation over diffusion. In the presence of curvature coupling due to protein-induced spontaneous curvature, these dynamics are altered and depend strongly on the strength of the spontaneous curvature induced by these proteins. These altered dynamics can be summarized as follows: for certain regimes of *Ŝ* and 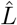, microdomains of proteins form on the membrane and are closely tied to the membrane curvature as is expected, generating a strong feedback between curvature and aggregation. We also found that for certain regimes of *Ŝ* and 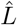, the growth rate decays, preventing the formation of protein aggregates and the membrane remains flat.

The interaction between curvature and protein aggregation in membranes has been studied in multiple modeling [21, 23, 34, 35], simulation [6, 7, 10], and experimental contexts [36–40]. Our work builds on this literature with a few key differences. Many of the theoretical models analyze the governing equations in simplified settings. In some cases, the geometry is fixed and the emergence of patterns is analyzed, and in other cases, the dynamics of the protein interactions on the surface is ignored [10, 23]. Here, we have analyzed the fully coupled system without any assumptions on the dominant regimes and demonstrated how curvature generation can affect aggregation. Another important feature of our model is the calculation of membrane tension. Since the lipid bilayer is assumed to be incompressible, the calculation of the Lagrange multiplier, which is widely interpreted as membrane tension (see detailed discussion in [30] and references therein), is an important aspect of the coupled physics. By incorporating the viscous nature of the membrane, we ensure that the incompressibility constraint is met rigorously at all times and therefore obtain the tension fields on the membrane. Our calculation of the heterogeneous tension fields are consistent with previous models as noted above and with experimental observations [41].

Finally, we discuss the relevance of our model in the context of experiments. The interaction between curvature and protein aggregation has been discussed in different membrane-protein systems and often these results are interpreted in the specific context of the particular experimental system [42–45]. These observations can lead to system-specific claims of whether membrane protein interactions result in a mechanochemical feedback between curvature and aggregation. In the context of specific biological processes such as endocytosis, aggregation of domains of protein-induced curvature is often assumed *a priori* or curvature is proposed as an organizing factor to explain cellular observations and experiments in reconstituted systems [46–52]. By developing a general theoretical framework that accounts for the coupled effects of protein diffusion, aggregation, and curvature generation, we have eliminated the need for such strong assumptions and more importantly, demonstrated that the intricate interactions between these different physics can lead to different regimes of pattern formation and membrane deformations. These regimes can be tuned and controlled by different parameters, allowing for exquisite control of experimental design. In summary, the comprehensive model that we have developed here allows for a broader interpretation and understanding of membrane-protein interactions in a unifying framework.

## 6 Acknowledgments

This work was supported by NIH NIGMS R01-132106 to P.R. and NSF CBET-1705377 to D.S.

## A Supplementary Figures

**Figure A.1:**
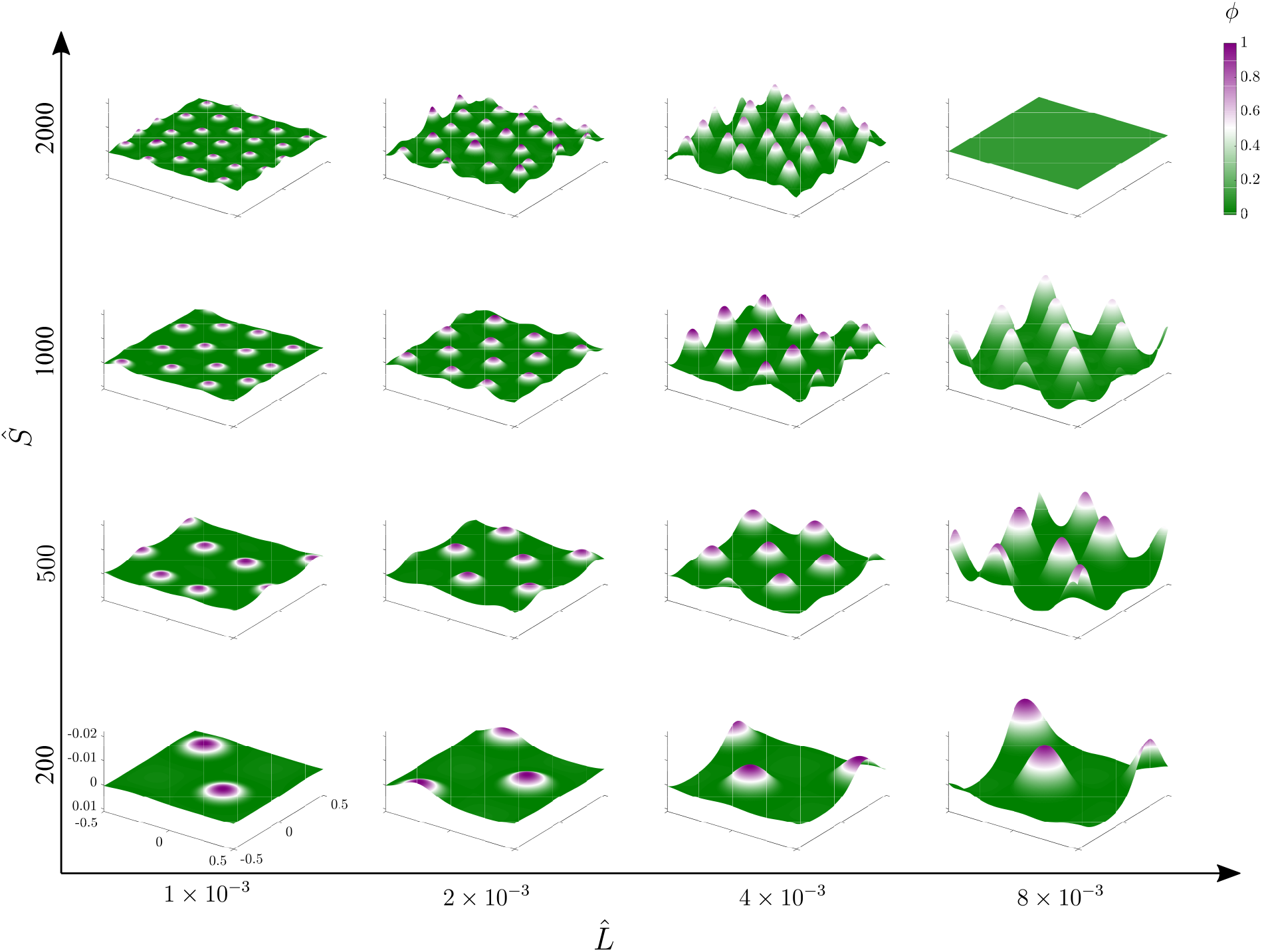
Protein distribution on the deformed membrane at a long time mimicking the steady state in the plane of 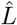 and *Ŝ*, with *Â* = 25.

**Figure A.2:**
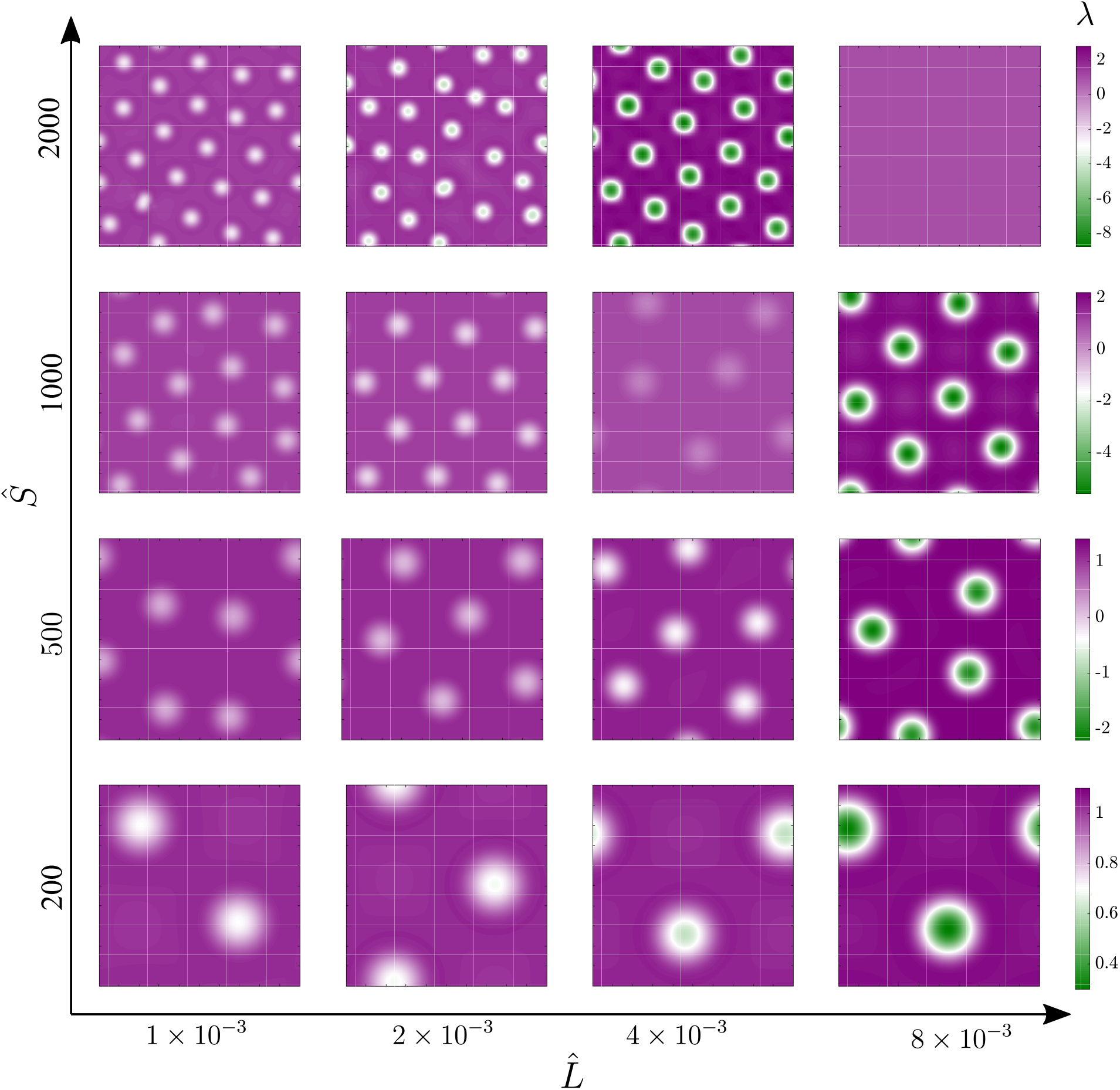
Membrane tension on the projected membrane surface at a long time mimicking the steady state in the plane of 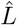 and *Ŝ*, with *Â* = 25.

## B Supplementary Movies

**[Movie M1](https://drive.google.com/file/d/1h-2WZgW5tpKKJssts3howy3HecwslYMH/view):** Time evolution of the protein density on a flat square membrane of area 1 *μ*m for *Â* = 25 and *Ŝ* = 200 in the Cahn-Hilliard case.

**[Movie M2](https://drive.google.com/file/d/19jSHyU0_hfL6UMa6_ZddD-9Sk-I7IpvN/view?usp=sharing):** Time evolution of the protein density on a flat square membrane of area 1 *μ*m for *Â* = 25 and *Ŝ* = 500 in the Cahn-Hilliard case.

**[Movie M3](https://drive.google.com/file/d/10WcSacJUYPI2ThyplGgtQOk8QNOt6PVB/view?usp=sharing)** Time evolution of the protein density on a flat square membrane of area 1 *μ*m for *Â* = 25 and *Ŝ* = 1000 in the Cahn-Hilliard case.

**[Movie M4](https://drive.google.com/file/d/1PLMvH0R5-PxBw7Ov7q5pnHyhfq5Zt46k/view?usp=sharing):** Time evolution of the membrane deformation and protein density in a square membrane of size 1 *μ*m^2^ for *Â* = 25, *Ŝ* = 200 and 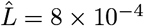 for the fully coupled system.

**[Movie M5](https://drive.google.com/file/d/1yb3EMkPSe5MNyMZVu6-DaTHH5Y_XJTvo/view?usp=sharing):** Time evolution of the membrane deformation and protein density in a square membrane of size 1 *μ*m^2^ for *Â* = 25, *Ŝ* = 1000 and 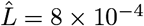 for the fully coupled system.

**[Movie M6](https://drive.google.com/file/d/18wFHeXzLyhrQ1ZL7FG0O02CzZrO7_OJl/view?usp=sharing):** Time evolution of the membrane deformation and protein density in a square membrane of size 1 *μ*m^2^ for *Â* = 25, *Ŝ* = 2000 and 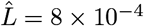 for the fully coupled system.

